# Phase separation of toxic dipeptide repeat proteins related to C9orf72 ALS/FTD

**DOI:** 10.1101/2020.01.31.926428

**Authors:** Hamidreza Jafarinia, Erik van der Giessen, Patrick R. Onck

## Abstract

The expansion mutation in the C9orf72 gene is the most common known genetic cause for amyotrophic lateral sclerosis (ALS) and frontotemporal dementia (FTD). This mutation can produce five dipeptide repeat proteins (DPRs), of which three are known to be toxic: poly-PR, poly-GR, and poly-GA. The toxicity of poly-GA is attributed to its aggregation in the cytoplasm, while for poly-PR and poly-GR several toxicity pathways have been proposed. The toxicity of the DPRs has been shown to depend on their length, but the underlying molecular mechanism of this length dependence is not well understood. To address this, a one-bead-per-amino-acid (1BPA) coarse-grained molecular dynamics model is used to study the single-molecule and phase separation properties of the DPRs. We find a strong dependence of the phase separation behavior on both DPR length and concentration, with longer DPRs having a higher propensity to phase separate and form condensed phases with higher concentrations. The critical lengths required for phase separation (25 for poly-PR and 50 for poly-GA) are comparable to the toxicity threshold limit of 30 repeats found for the expansion mutation in patient cells, suggesting that phase separation could play an important role in DPR toxicity.

## Introduction

Hexanucleotide repeat expansion G_4_C_2_ in the C9orf72 gene is the most common genetic mutation in familial cases of amyotrophic lateral sclerosis (ALS) and frontotemporal dementia (FTD) (1, 2). Healthy individuals typically have less than around 20 repeats of this expansion, while in most patient cells the size of the expansions is estimated to be between several hundred and several thousand repeat units (1–3). There is no consensus on the critical expansion size for the onset of the disease and different cut-offs between 30-80 repeats have been reported for the toxicity threshold (1, 2, 4, 5).

The pathology initiated by the repeat expansion has been proposed to affect a wide range of cellular processes (6). The three main mechanisms of toxicity are: loss of function of C9orf72 proteins (1, 7), and toxic gain of function from the repeat expansion itself (8, 9) or from dipetide repeat proteins (DPRs) translated from sense and antisense transcripts of the repeat expansion (10–12). It has been shown that DPRs are capable of inducing toxicity without the repeat expansion in different cell types (12–17).

Repeat-Associated Non-AUG (RAN) translation of the sense and antisense transcripts of the repeat expansion from all reading frames can produce five types of DPRs: poly-PR, poly-GR, poly-GA, poly-GP, and poly-PA (10, 11). Poly-PR, poly-GR and poly-GA can induce length-dependent and dosage-dependent toxicity (12, 14, 16, 18–21) of which especially the R-DPRs, i.e. poly-PR and poly-GR, are highly toxic. Poly-PR is known to be the most toxic DPR (12–15, 20, 22). Several studies indicate no significant toxicity for poly-GP and poly-PA (12, 20, 23). In the current study we use the term toxic DPRs to refer to poly-PR, poly-GR and poly-GA.

Poly-GA is the most abundant and the most aggregation prone DPR (10, 22). Poly-GA has only a few interactors in the cell (22). It has been suggested that poly-GA toxicity is due to the formation of cytoplasmic aggregates and direct sequestration of proteins (24–28). Poly-GA aggregates are shown to impair the nuclear import of TDP-43 (29) and enhance DNA damage (27). Poly-PR and Poly-GR have many target proteins inside the cell (22, 30) and are likely to be involved in several pathology pathways (reviewed in (6)). Significant attention has been drawn to nucleocytoplasmic transport defects, changes in the dynamics of membrane-less organelles through impaired liquid-liquid phase separation (LLPS), and nucleolar dysfunction (13, 14, 16, 17, 22, 31, 32). Recently, the toxicity of Poly-PR has been related to changes in LLPS of heterochromatin protein 1*α* (HP1*α*) (17) and nucleophasmin (NPM1) (16) through their direct interaction with poly-PR inside the cell nucleus.

Depending on the type of DPR and the toxicity mechanism, the DPR pathology can start either in the cytoplasm, nuclear pore complex (NPC) or nucleus. Despite the recent progress, a clear understanding is lacking on how the DPRs cause neurotoxicity in C9orf72 ALS/FTD. Due to the methodological difficulty of synthesizing repetitive sequences, it is highly challenging to study the length-dependent properties of DPRs. As a result, almost all previous *in vitro* cell-free studies were limited to DPRs with less than 30 repeats (21, 22, 33, 34) (only one recent study used 60 repeats of poly-PR (16)), which might not mimic the exact role of long DPRs in patients (35). Moreover, the concentration inside the condensed phases, which is important for further maturation of the phase separated condensates (36), are not easily determined in experiments. These problems can be overcome by using experimentally-calibrated coarse-grained molecular dynamics (MD) models that can capture the sequence specificity and are suitable for simulating high-density phases of proteins (37–39). In this study we use our one-bead-per-amino-acid (1BPA) coarse-grained MD model (37, 40, 41) to investigate the single-molecule and phase separation behavior of toxic DPRs in an attempt to identify possible mechanistic roles of DPRs in generating toxicity.

## Results and Discussion

### Single-molecule properties of toxic DPRs

The dynamics of intrinsically disordered proteins (IDPs) is crucial for their function, including, for example, recognition and binding to target molecules (43–45). The dimension of IDPs can have a large effect on their functional properties. At a fixed temperature and solvent condition the dimension of an IDP is determined by its amino acid sequence (42, 43). The repetitive sequences of toxic DPRs contain only two types of amino acids (see Fig. 1*A*). Therefore, the overall properties of the DPRs strongly depend on the physiochemical properties of Glycine, Alanine, Proline, and Arginine, and possibly the patterning of these residues. Alanine is hydrophobic, Arginine is positively charged, Proline contributes to the rigidity and Glycine to the flexibility of the protein backbone (46–48).

**Fig. 1.**
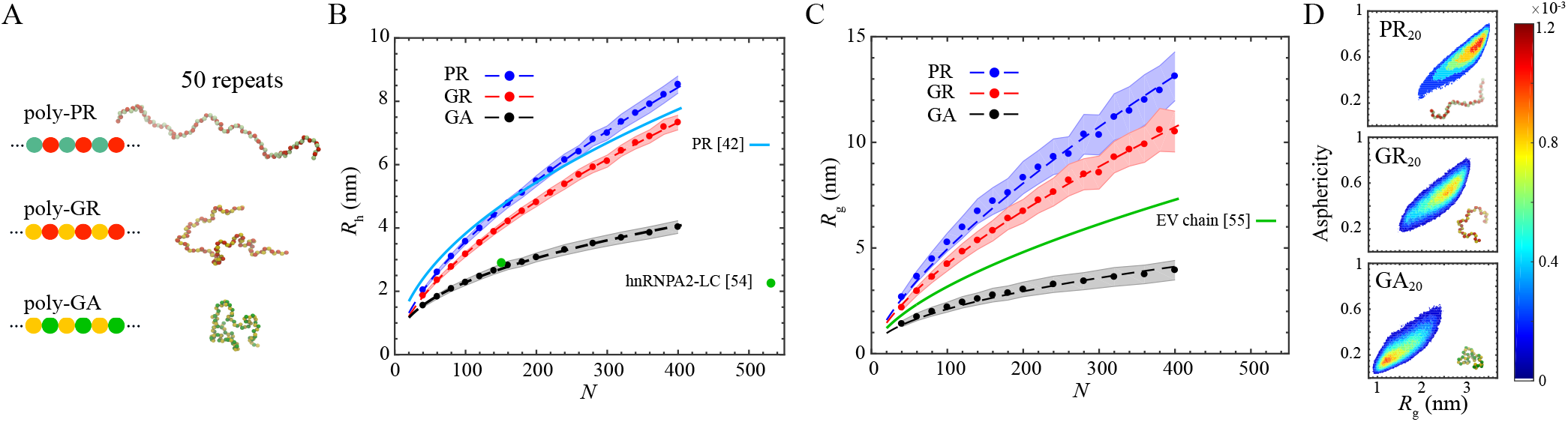
Comparison between the single-molecule properties of poly-PR, poly-GR, and poly-GA. (A) Schematic representation of the DPRs (left) and simulation snapshots (right). (B) Hydrodynamic radius *R*_h_ of the DPRs plotted against their chain length *N* (circles fitted with dashed lines). The solid light-blue line is a prediction for the *R*_h_ of poly-PR based on a closed-form expression (42). Experimental *R*_h_ value for the low-complexity (LC) domain of hnRNPA2 is shown by a green circle. (C) Radius of gyration *R*_g_ of the DPRs plotted against their chain length *N* (circles fitted with dashed lines). The *R*_g_ of an excluded volume (EV) chain is provided for comparison. In (B) and (C) the shaded regions indicate half of the standard deviation. (D) Two-dimensional probability distribution of the Asphericity and *R*_g_ for PR_20_, GR_20_, and GA_20_.

We use our one-bead-per-amino-acid (1BPA) molecular dynamics model (details are provided in *SI Appendix*) to study the dimensions of the three toxic DPRs. In Fig. 1*B* and *C* we show the hydrodynamic radius, *R*_h_, and the radius of gyration, *R*_g_, of poly-PR, poly-GR, and poly-GA. The results are shown for a range of chain lengths *N* between 40−400, with *N* being two times the number of repeats, *n*. For the same chain length, the dimension of poly-PR is larger than Poly-GR which is larger than poly-GA. The electrostatic repulsion between the uniformly distributed Arginine residues results in more expanded conformations for the R-DPRs compared to the hydrophobic poly-GA. This observation is consistent with previous studies on the effect of charged residues and their patterning on the dimension of IDPs (49–51). The observed difference between poly-PR and poly-GR, however, cannot be explained by the non-bonded interactions alone. Proline is more hydrophobic than Glycine (37, 52) and thus poly-PR is expected to have a more compact structure than poly-GR. Therefore, the larger *R*_h_ and *R*_g_ values for Poly-PR can only be attributed to the different contribution of Proline and Glycine to the backbone stiffness. Indeed, Proline is much stiffer than Glycine due to the cyclic structure of its side chain (46–48), which is incorporated in the 1BPA force field (53). Our results are consistent with the observed correlation between Proline content and extended conformation of IDPs (42, 47).

In Fig 1*B* we compare our simulation results for *R*_h_ of poly-PR with the Marsh and Forman-Kay fit to the experimental *R*_h_ values of 36 IDPs (42). The suggested expression takes into account the Proline content and the absolute net charge of the chain. The difference between our simulation results and the prediction by Marsh and Forman-Kay expression is less than 16%. The observed difference for longer chains can be due to the fact that the patterning of amino acids has not been considered as an input variable in the suggested expression (42). To show the importance of sequence patterning, for instance, the *R*_h_ of three variants of Proline-Arginine chains with the same amino acid composition but different patterning of Proline and Arginine residues are depicted in *SI Appendix* Fig. S2. These results show that the chain favors a conformation with the highest *R*_h_, when the Proline and Arginine residues are well mixed, as in poly-PR. Poly-GA forms the most compact conformation due to the uniform distribution of hydrophobic Alanine residues and the low stiffness of the Glycine residues. We also compare the simulated *R*_h_ of poly-GA with the experimental *R*_h_ of the disordered low-complexity (LC) domain of hnRNPA2 (54), see Fig. 1*B*. The hnRNPA2 LC domain contains hydrophobic residues (mainly Phenylalanine and Tyrosine) distributed along the sequence. It has been suggested that the high Glycine content (47%) of hnRNPA2 LC domain contributes to its compactness (54). With the same chain length the hydrodynamic size of poly-GA is very similar to that of the hnRNPA2 LC domain.

Relating the *R*_g_ of the DPRs to the chain length *N* via *R*_g_ ∝ *N*^*ν*^ (55) leads to scaling exponents of *ν* = 0.70±0.02 for poly-PR, 0.67±0.02 for poly-GR, and 0.48±0.02 for poly-GA. Similar scaling exponents of around 0.7 have been obtained for extended variants of prothymosin *α* (ProT*α*) (with a mean net charge per residue of −0.46) in water (55). The scaling exponent of poly-GA is close to the value *ν* = 0.5 expected for a random coil, i.e., a polymer in a theta solvent. The *R*_g_ ∝ *N*^0.6^ of an excluded volume (EV) chain is also plotted for comparison in Fig. 1*C* (55). Poly-PR and poly-GR have an *R*_g_ that is larger than an EV chain due to the repulsion of like charges, while the flexible and more hydrophobic poly-GA has a lower *R*_g_. In Fig. 1*D* we compare the asphericity, which measures the chain shape, vs. *R*_g_ plots for DPRs with a repeat length *n* = 20. PR_20_ samples conformations with a larger *R*_g_ and asphericity than GR_20_ and GA_20_, showing that poly-PR is more extended and assumes shapes closer to a rod-like conformation.

### Poly-GA phase separation

Poly-GA forms cytoplasmic aggregates (5, 18, 22, 25) that are relatively stable in photobleaching experiments (22). Fourier Transform Infrared Spectroscopy (FTIR) measurements show a random coil structure for GA_15_ right after incubation (21). After a certain incubation time, the GA_15_ molecules form aggregates as indicated by a change in the average particle size in the system (21). After a few hours, GA_15_ starts to form fibrils containing cross-*β* sheet structures with disordered molecules of poly-GA still in solution (21, 56).

Our results show that poly-GA undergoes a length-dependent phase separation to form a condensed (high-density) phase and a dilute (low-density) phase (Fig. 2). The condensed phases of poly-GA are spherical and exchange molecules with the surrounding (see snapshots in Fig. 2*A*, *SI Appendix*, Fig. S3). Since hydrogen bonding is not included in the modeling, we are not able to predict the final transition into relatively stable aggregates or higher-order *β*-type structures of poly-GA as observed in the experiments (21, 22, 25). However, our results do suggest that long-range hydrophobic interactions drive the formation of high-density phases of Poly-GA, which brings the residues close enough for short range hydrogen bonds to form (57). Further transition of liquid condensates to more solid-like structures have been experimentally observed for FUS and hnRNAPA1 (58–60). To find the critical repeat length required for phase separation we constructed the coexistence phase diagram of poly-GA (Fig. 2*A*) by determining the concentrations of the condensed phase (*ρ*_H_) and dilute phase (*ρ*_L_) using the slab method (Fig. 2*B*) (38, 61, 62). In the phase diagram in Fig. 2*A* the vertical axis is the repeat length *n* and the horizontal axis is the concentration *ρ*. Outside the coexistence curve (i.e. for concentrations lower than *ρ*_L_, higher than *ρ*_H_ and lengths shorter than the critical repeat length) there is only a uniform phase, while inside it, the poly-GA molecules phase separate (see the snapshots at the left in Fig. 2*A*). The black arrows in Fig. 2*A* show the concentrations of the two phases. The critical repeat length for phase separation of poly-GA is found to be *n* = 50. Below this critical repeat length no phase separation is observed at any concentration. This critical repeat length is in good agreement with the critical range of 46 < *n* < 61 found by Yamakawa et al. for the formation of aggregates of poly-GA in Neuro2a cells (5).

**Fig. 2.**
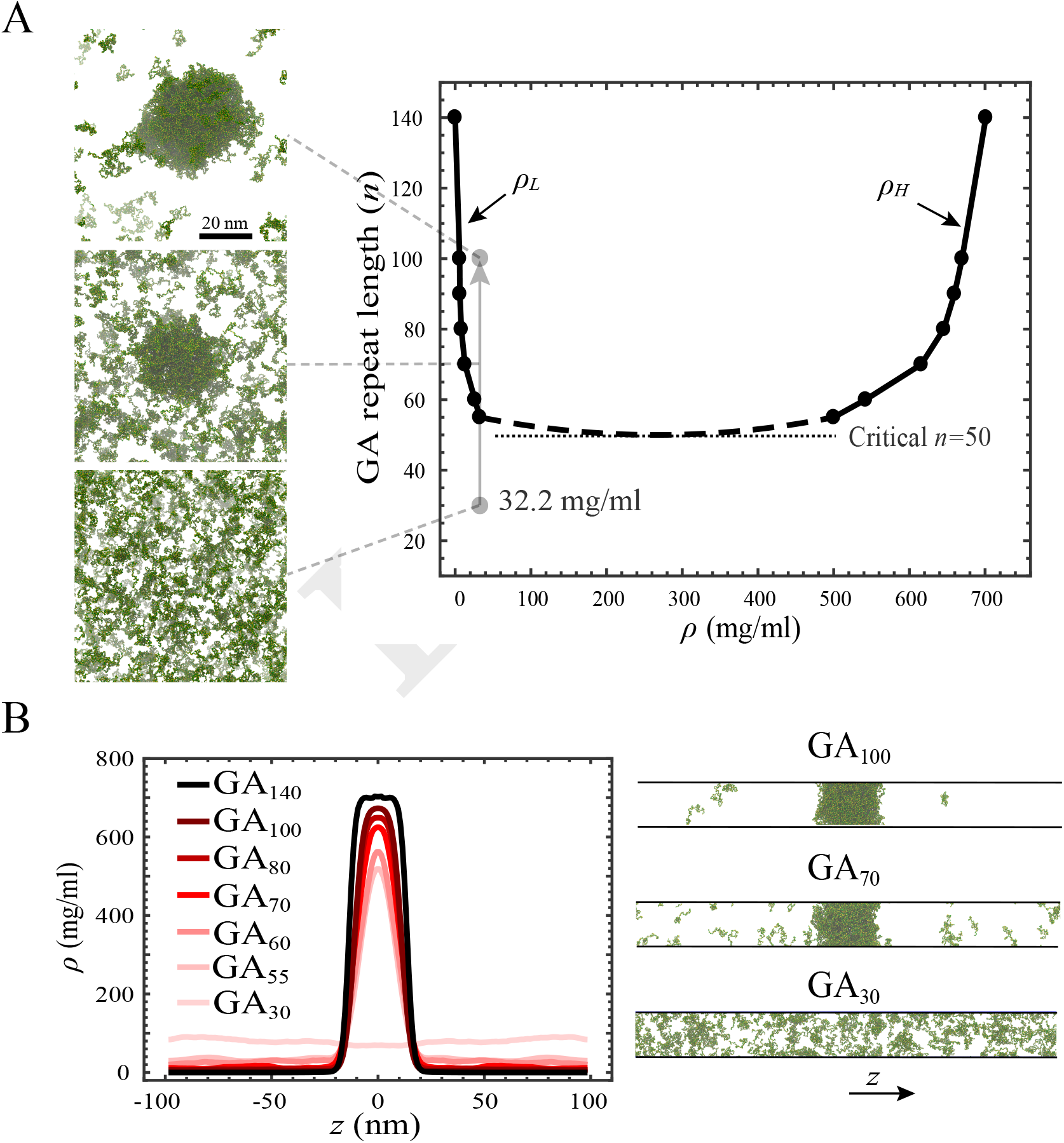
Poly-GA length-dependent phase separation and coexistence phase diagram. (A) Coexistence phase diagram of poly-GA obtained from slab simulations. Phase separation occurs above the critical repeat length *n* = 50 inside the coexistence region. *ρ*_H_ and *ρ*_L_ are the concentrations of the condensed phase and dilute phase, respectively. Simulation snapshots at the left show the results for GA_100_, GA_70_, and GA_30_ for a total concentration of 32.2 mg/ml at equilibrium. (B) Poly-GA slab density profiles (left panel) and sample slab simulation snapshots (right panel).

The values of *ρ*_H_ and *ρ*_L_ depend on the repeat length, with longer poly-GA molecules forming condensed phases with higher concentrations and dilute phases with lower concentrations (see Fig. 2*A*). The same trend has been observed in previous simulation studies for polymers and IDPs with different lengths (38, 61). Also in experiments using synthetic molecules the saturation concentration, which is equal to *ρ*_L_ at equilibrium, was observed to decrease with chain length (63). For 60 ≤ *n* ≤ 100 we find the concentration range to be 6−33 mg/ml for the dilute phase, and 500-670 mg/ml for the condensed phase. Using linear interpolation in Fig. 2*A* the molar concentration of the GA_76_ condensed phases is found to be ≈ 34 mM, comparable to that of the 151 residue long hnRNPA2 LC domain (30-40 mM) obtained in experiment (54). This is expected since the hnRNPA2 LC domain has a similar dimension as GA_76_ (see Fig. 1*B*).

Cluster size distribution (CSD) analysis of poly-GA at equilibrium shows a strong correlation between the poly-GA total concentration and its ability to phase separate at small concentrations at the left of the phase diagram (*SI Appendix*, Fig. S4 and Fig. 2*A*). When phase separation occurs, a further increase of the total concentration has no effect on the *ρ*_H_ and *ρ*_L_, but only increases the size of the condensed phase (*SI Appendix*, Fig. S5). The CSD results of *SI Appendix*, Fig. S4 also reveal, see *SI Appendix*, Fig. S6, that at a fixed concentration, the number of free molecules *N*_free_ is higher for shorter dipeptides and that, as expected, a drop in *N*_free_ can be taken as an indication of phase separation.

### Phase separation of R-DPRs

R-DPRs have been observed to form liquid droplets in the presence of RNA (33, 34), phosphate ions (33, 56) and several RNA-binding proteins (16, 22). Previous studies have pointed at the important role of electrostatic and cation-*π* interactions in the LLPS of R-DPRs with multivalent proteins and RNA molecules for these cases. (16, 22, 33, 34). The phase separation of PR_30_ with different polyanions, known as complex coacervation, has recently been investigated using both *in vitro* experiments and coarse-grained Dynamic Monte Carlo (DMC) modeling (34).

Our simulations show no phase separation of poly-PR and poly-GR, as a consequence of the electrostatic repulsion between Arginine residues, see *SI Appendix*, Fig. S7 (left panels). The same result has been obtained in *in vitro* experiments with PR_20_ in the presence of monovalent ions (33). The addition of anionic homopolymers (acidic molecules) of poly-Aspartate (poly-D) to the simulation box induce the phase separation of R-DPRs into liquid droplets, *SI Appendix*, Fig. S7 (right panels). R-DPRs bind to acidic molecules, and upon binding they become more compact (see Fig. 3*A*). In *SI Appendix*, Fig. S8*A* we compare the potential of mean force (PMF) associated with binding of the R-DPRs, with repeat lengths *n* = 30, to an acidic molecule of length *N* = 60. The binding of poly-GR to the acidic molecule is stronger than poly-PR as the flexibility of poly-GR allows it to make more number of contacts with an acidic molecule than poly-PR, see *SI Appendix*, Fig. S8*B*.

**Fig. 3.**
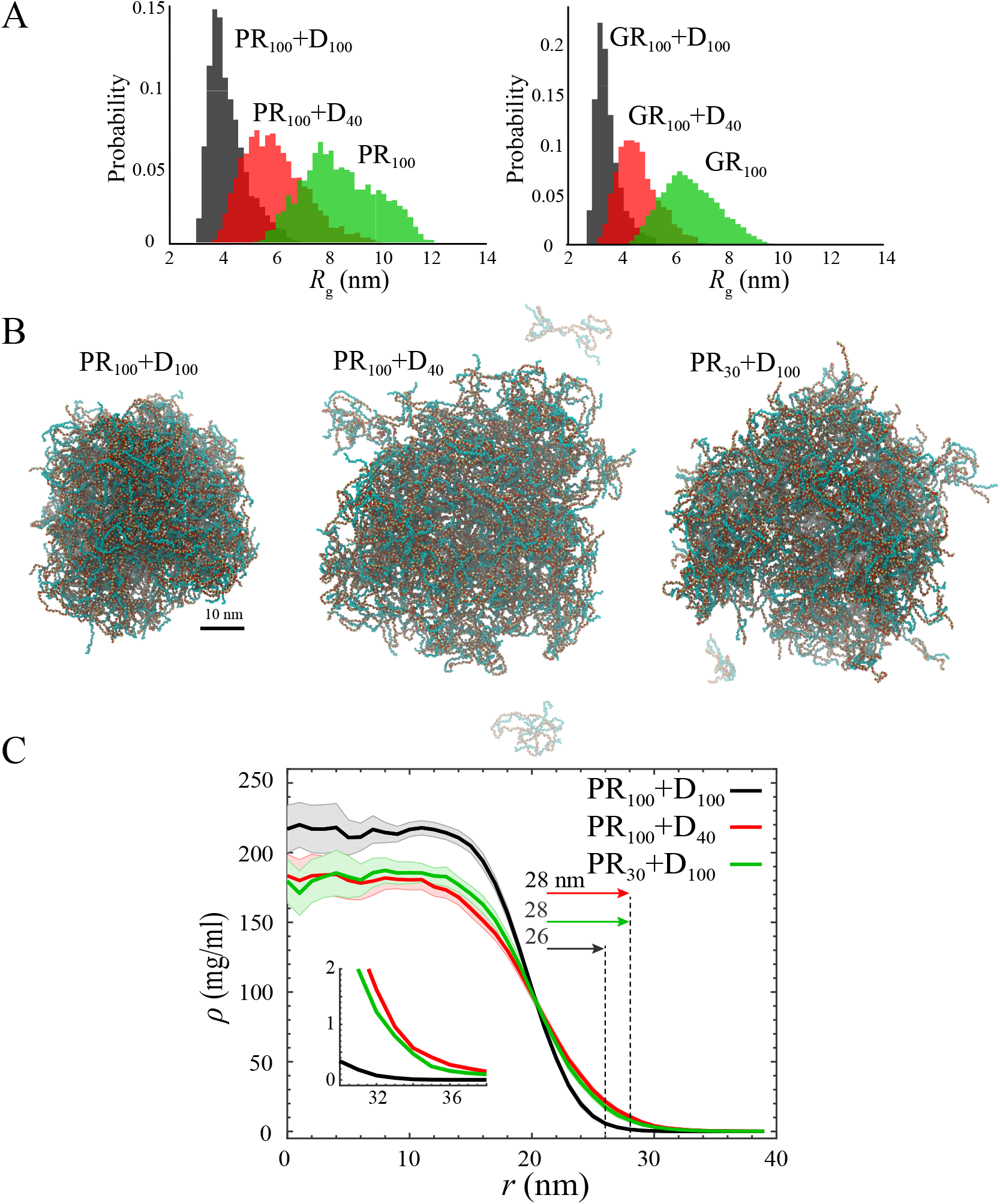
Binding of R-DPRs to acidic molecules (poly-D) and length-dependent droplet formation of poly-PR. (A) Histograms for *R*_g_ of single PR_100_ (left) and GR_100_ (right) in the presence and absence of a D_40_ and D_100_ molecule. Binding to acidic molecules reduces the *R*_g_ of R-DPRs with longer acidic molecules having a larger effect. (B) Simulation snapshots for length-dependent droplet formation of poly-PR with acidic molecules for a total concentration of 14.8 mg/ml and a poly-PR concentration ratio of *r*_PR_ = 0.57. (C) Radial density profiles measured from the center of mass of the droplets presented in (B). Light shades indicate half of the standard deviation as error bars. The vertical dashed lines indicate the droplet size (see the details for droplet radius calculation in the *SI Appendix*). The inset shows the zoomed density profiles for *r* > 30 nm.

Long acidic tracts with length ranges of 12-41 amino acids can be found in the disordered regions of two nucleolar proteins: Nucleolin (NCL_1−300_) and Nucleophasmin (NPM1_120−240_), see the net charge per residue (NCPR) histograms in *SI Appendix*, Fig. S9 (top panels). NCL mislocalization and disruption has been observed in ALS patient cells (64). NSR1, the yeast homolog of NCL, has also been observed to be a strong modifier of PR_50_ toxicity. The toxicity was shown to be suppressed by deletion and enhanced by upregulation of NSR1 (14). A recent study also suggested that poly-PR-induced NPM1 mislocalization generates toxicity (16). Our simulations show binding of the R-DPRs to both NPM1_120−240_ and NCL_1−300_ through interaction with the acidic tracts, *SI Appendix*, Fig. S9 (bottom panels). These results are consistent with the experimentally observed binding of full-length NPM1 and NCL to the R-DPRs (16, 22).

Fig. 3*B* and *C* and Fig. 4 show that the phase separation of poly-PR not only depends on its length but also on the length of the acidic molecules. At a fixed total mass concentration, and mass concentration ratio of poly-PR, *r*_PR_ = (poly-PR mass concentration)*/*(total mass concentration), longer repeats of poly-PR and poly-D phase separate to smaller droplets with a higher concentration surrounded by a dilute phase with a lower concentration (see Fig. 3*C*). Reduction of the length of acidic molecules (Fig. 3*B* middle panel) or poly-PR (Fig. 3*B* right panel) increases the size of the droplet and the concentration of the dilute phase, see Fig. 3*C*.

**Fig. 4.**
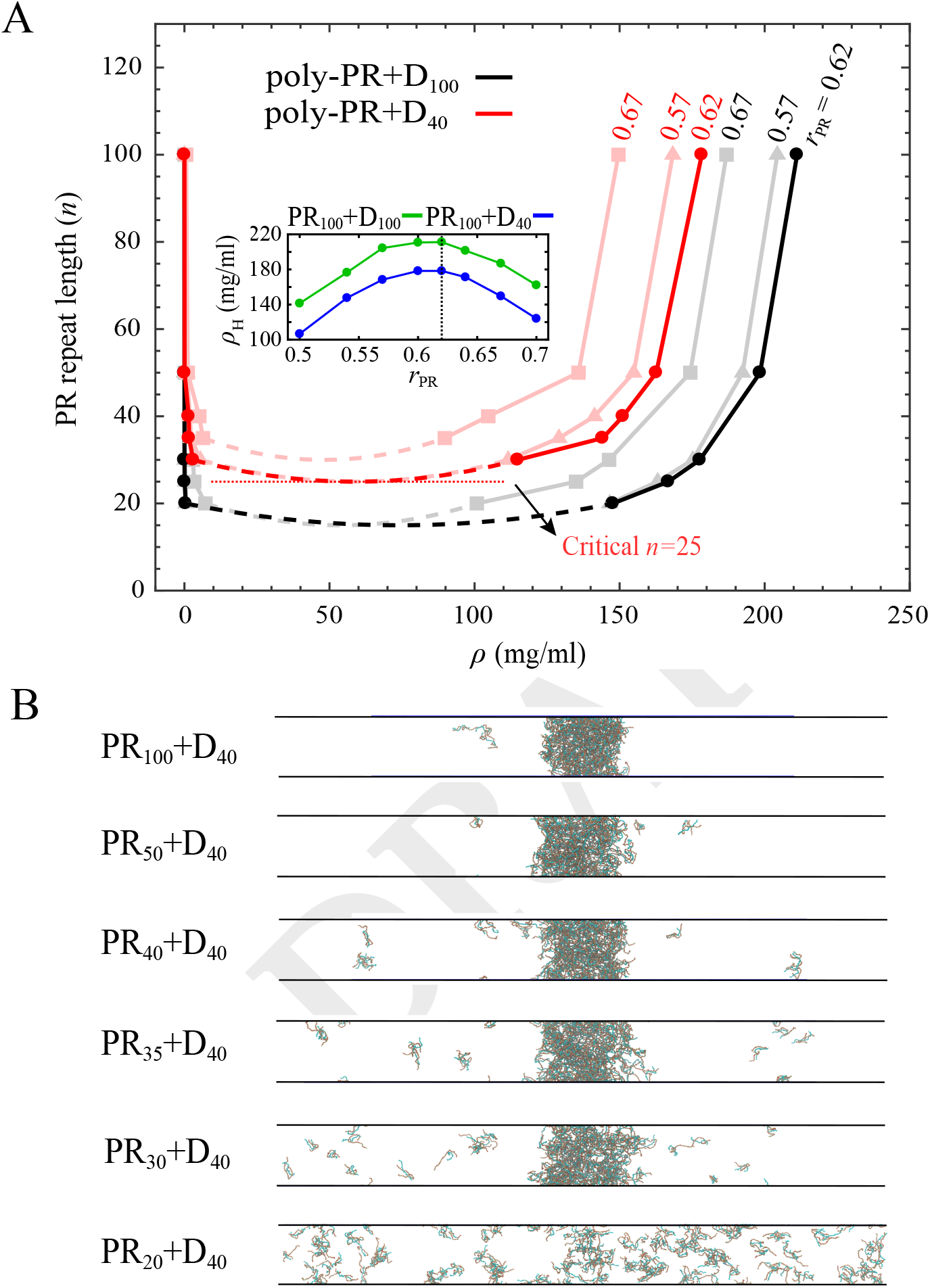
Length-dependent phase separation of poly-PR. (A) Phase diagrams for poly-PR phase separation with acidic molecules of lengths 40 and 100 for different Poly-PR concentration ratios of *r*_PR_ = 0.57, 0.62, and 0.67, see the tilted numbers next to each phase diagram. Light shades are used for the phase diagrams constructed for *r*_PR_ = 0.57 and 0.67. The minimum critical repeat length for phase separation of poly-PR with D40 is indicated with a horizontal dotted line. The inset shows the concentration of the high-density phases of PR_100_ + D_40*/*100_ for different poly-PR concentration ratios. (B) Slab simulation snapshots for length-dependent phase separation of poly-PR with D40 for a poly-PR concentration ratio of *r*_PR_ = 0.62. No phase separation can be observed below a critical poly-PR repeat length *n* = 25.

To further study the phase separation of poly-PR, we use slab simulations to obtain the coexistence phase diagrams in Fig. 4*A*. The phase diagrams are constructed for acidic poly-D molecules of lengths *N* = 40 and 100 and three different poly-PR concentration ratios *r*_PR_ = 0.57, 0.62, and 0.67. Similar to the trend observed in Fig. 3*B* and *C*, we observe that longer repeats form condensed phases with higher concentrations and dilute phases with lower concentrations, in agreement with experimentally observed length-dependent complex coacervation of oppositely-charged polyelectrolytes (65). The poly-PR condensed phase concentration ranges from 90 to 211 mg/ml, which is much lower than the concentration range 500-670 mg/ml obtained for poly-GA for *n* ≤ 100. As can be seen in the phase diagram, the critical poly-PR repeat length required for phase separation is lower for longer acidic molecules. The results in Fig. 4*A* show an optimum concentration ratio of *r*_PR_ = 0.62 for phase separation of poly-PR with acidic molecules. At this concentration ratio, with fixed peptide lengths, poly-PR forms condensed phases with the highest concentration, see the inset of Fig. 4*A*. In Fig. 4*A* we constructed the phase diagrams for two other concentration ratios of *r*_PR_ = 0.57 and 0.67 around the optimum value. For lower (< 0.57) and higher (> 0.67) *r*_PR_ values the concentration of the condensed phase is smaller and the critical repeat length is larger. For *r*_PR_ > 0.8 and *r*_PR_ < 0.3 we observed no phase separation (data not shown). Our results are consistent with electrostatic-driven phase separation of R-DPRs with full-length NPM1, with phase separations that depend on concentration ratios of proteins and length of the R-DPRs (16, 22). For D_40_, which has almost the same length as the longest acidic tract in the nucleolar targets of R-DPRs (see *SI Appendix*, Fig. S9), the critical repeat length of poly-PR required for phase separation is found to be *n* = 25 for the optimum concentration ratio *r*_PR_ = 0.62 (see the phase diagram in Fig. 4*A* and the slab simulation snapshots in Fig. 4*B*). Below this critical repeat length, poly-PR can bind to acidic molecules but these small clusters are unable to phase separate, see Fig. 4*B*. The slab density profiles used to obtain the phase diagram are presented in *SI Appendix*, Fig. S10.

In *SI Appendix*, Fig. S11 we compare the coexistence phase diagram of poly-PR and poly-GR with D_100_ obtained for the same concentration ratios of *r*_PR_ = *r*_GR_ = 0.62. At a fixed repeat length *n*, poly-GR forms condensed phases with higher concentrations than poly-PR. This observation can be attributed to the sizes of the dipeptides and their free energy of binding to acidic molecules. With the same length, Poly-GR is more compact than poly-PR irrespective of the presence of acidic molecules (see Fig. 1 and Fig. 3*A*). This, together with the stronger binding of poly-GR to an acidic molecule (see *SI Appendix*, Fig. S8*A*), explains the differences in the phase diagrams. Previous experimental measurements have indicated greater association of NCL with poly-GR than poly-PR (22). Consistent with this observation, photobleaching experiments for a single nucleolus labeled by GFP-NCL have shown that poly-GR is more effective than poly-PR in reducing the mobile fraction and fluorescence recovery rate of GFP-NCL (22). Our results suggest that the more stable physical interaction of poly-GR with the nucleolar components can be attributed to the stronger binding of poly-GR to the acidic tracts inside the nucleolus.

## Conclusions

We showed in this work that poly-PR favors the most extended conformations among all three toxic DPRs due to the patterning of charged Arginine residues and the contribution of Proline residues to the backbone rigidity. Our findings showed a stronger binding of poly-GR to acidic molecules compared to poly-PR which can be attributed to the more flexible backbone of poly-GR enabling the chain to make more contacts with the acidic molecule. We observed that longer DPRs form condensed phases with higher concentrations and dilute phases with lower concentrations. For Poly-GA we found that increasing the concentration increases the propensity for phase separation at small concentrations and increases the size of the condensed phase at larger concentrations, both consistent with the expected trend for liquid condensates (66). Increasing the length of DPRs increases the number of possible interactions and results in an increase in the multivalency of the system. Previous experiment and simulation studies have shown that a critical number of valences is required for the formation of biomolecular condensates (39, 48, 63, 67, 68). This observation is in agreement with the phase diagrams presented in Fig. 2*A* and Fig. 4*A*. We observed an inverse correlation between the *ρ*_H_ and *R*_g_ of the DPRs. With the same chain lengths and the same concentration ratios of R-DPRs the *ρ*_H_ of poly-PR is lower than poly-GR which is lower than poly-GA, conceptually consistent with the observed correlation between the compactness of IDPs and their tendency to phase separate (69).

Our results for poly-GA repeat lengths larger than 50 suggest that aggregate nucleation starts with liquid phase separation as experimentally observed for several RNA-binding proteins (36, 58). Care should be taken in comparing our poly-GA droplets with the insoluble aggregates observed in previous studies (21, 22, 25) since our 1BPA force field does not take into account secondary or higher order structure formation of poly-GA. However, our coarse-grained MD model does capture the length-dependent aggregation of poly-GA (5). The critical repeat length of *n* = 50 for phase separation of poly-GA is in good agreement with the critical range of 46 < *n* < 61 found in Neuro2a cells for the formation of aggregates of poly-GA (5). The longest acidic tract found in the nucleolar targets of R-DPRs is 41 (see *SI Appendix*, Fig. S8). We found the minimum critical repeat length for the phase separation of the most toxic DPR, i.e. poly-PR, with similar-sized acidic chains D_40_ to be *n* = 25 which is close to the lower toxicity threshold limit of 30 repeats found for the C9orf72 expansion mutation in patient cells (1, 2, 4), suggesting that phase separation into liquid droplets could play an important role in DPR-mediated ALS/FTD.

## Materials and Methods

We use our coarse-grained 1BPA model (37, 53) that: 1-differentiates between the bonded potentials of Glycine, Proline, and other residues, and 2-is fine-tuned to capture the properties of poly-Proline, poly-Glycine and FG-Nup segments with the highest Arginine content. More details are provided in the *SI Appendix*. Langevin dynamics simulations are performed at 300 K and physiological salt concentration of 150 mM using GROMACS. Droplet simulations are performed in a cubic box of size 80 nm. Slab simulations are conducted based on the procedure described in (38). All simulations are performed for t ≈ 3 *μ*s which is sufficient to obtain converged density profiles. The density profiles are calculated using discrete cells of thickness 1 nm and time-averaged for at least 1 *μ*s at equilibrium. The critical repeat lengths required for phase separation are obtained with an error of less than 5 repeats. More details are provided in the *SI Appendix*.

## ACKNOWLEDGEMENTS

We thank Liesbeth Veenhoff for helpful discussions and Anton Jansen for assistance in calculating the hydrodynamic size of the proteins. We are also grateful for use of the computational cluster Peregrine of the University of Groningen.

