## Supporting_information for "Phase separation of toxic dipeptide repeat proteins related to C9orf72 ALS/FTD"

DRAFT

### Coarse-grained force field

**Update of the 1BPA coarse-grained force field.** The 1BPA force field has been used before to study intrinsically disordered FG-Nups and nucleocytoplasmic transport (1–3). The bonded interactions, i.e. bending and torsion potentials, in this force field are residue and sequence specific. This force field, interestingly, differentiates between the bending and torsion potentials of Glycine, Proline, and other residues. This feature is highly important since DPRs are rich in Proline and Glycine and it has been shown that these two residues contribute to the rigidity and flexibility of an IDP (4–7). Therefore, using the 1BPA model enables us to distinguish between the properties of poly-PR and poly-GR and to obtain more accurate results for poly-GA. The non-bonded interactions in this force field have been calibrated against the experimentally known $R_{\mathrm{h}}$ values of FG-Nup segments (8). For Proline and Glycine the interaction parameters have also been fine-tuned using the end-to-end distance and radius of gyration of poly-Proline (9) and poly-Glycine segments (10) which makes the 1BPA a proper choice for investigating the properties of DPRs.

The majority of FG-Nup segments, however, contain less than $0.6\%$ of Arginine (R) (see the pink shaded band in Fig. S1). For the ones with more than $0.6\%$ of R a correlation can be observed between the R content and the $R_{\mathrm{h}}$ error (see black dashed line in Fig. S1) showing that there is still room for improving the R interaction with other residues in the 1BPA force field. To achieve this we further fine-tune the relative hydrophobic strength ($\varepsilon_i$) of R. The $\varepsilon_i$ value is a residue-specific parameter that ranges between 0 and 1 and is close to 0 for hydrophilic polar residues. Since we also study the interaction of R-DPRs with acidic molecules, we recalibrate the $\varepsilon_i$ values of all charged residues, i.e. RDEK. The aim is to obtain an updated 1BPA force field that is more accurate for studying the properties of R-DPRs and their interaction with negatively-charged molecules.

To update the 1BPA force field we slightly increase the $\varepsilon_i$ values of charged residues to 0.005 (see Table S1), thus reducing the $R_{\mathrm{h}}$ error for all the six FG-Nup segments with R content $> 0.6\%$ (see Fig. S1$A$). This choice of parameter for the relative hydrophobic strength of charged residues gives the best results in terms of the total average error and the minimum largest error in our calibration simulations with 16 FG-Nup segments presented here and originally used for the calibration of 1BPA (2). The total average and the largest errors are found to be $8.3\%$ and $21.1\%$ in the 1BPA force field, and $7.5\%$ and $17.1\%$ in the updated 1BPA force field. The correlation mentioned earlier for an R content $> 0.6\%$ still exists in the updated 1BPA (red dashed line) which might be due to the absence of cation-$\pi$ interactions between R and residues with aromatic rings in our force field. However, this has no effect on our simulation results since the DPRs and acidic molecules studied in this work contain no aromatic residues. A direct comparison between the two force fields is presented in Fig. S1$B$. At physiological intracellular pH between 7 and 7.4, i.e more than three pH units away from the p$K_{\mathrm{a}}$ values of Arginine and Aspartic acid, we assume R, D, E, and K to be fully charged (11).

**Complex coacervation of R-DPRs.** The complex coacervation of polyelectrolytes is driven by a combination of enthalpic and entropic effects (12). Coulombic energy change and counterion release entropy are the main contributors to the free energy of complexation (13). In our single-molecule and phase separation simulations of R-DPRs with stretches of acidic amino acids, we account for the screening effect of ions, but similar to previous theoretical (14, 15) and coarse-grained MD models (16, 17) used to study complex coacervation, the effect of counterion condensation has not been considered in our modeling. Despite this limitation, our simulations capture the experimentally observed length-dependence of $\rho_{\mathrm{L}}$ (concentration of the dilute phase) and $\rho_{\mathrm{H}}$ (concentration of the condensed phase) for polyelectrolytes (18). Here we compare the effect of Coulomb energy change and counterion release entropy for the complexation of R-DPRs with acidic molecules by calculating the Coulomb strength parameter as suggested by Ou and Muthukumar (13). In their study the Coulomb strength parameter $\Gamma$ has been defined as

$$\Gamma = \frac{l_{\mathrm{B}}}{l_0}.$$

Here $l_{\mathrm{B}} = e^2/(4\pi\epsilon_0\epsilon_r k_{\mathrm{B}}T)$ is the Bjerrum length, where $e$ is the elementary charge, $\epsilon_0$ is the vacuum permitivity, $\epsilon_r$ is the relative permitivity, $k_{\mathrm{B}}$ is the Boltzmann constant, $T$ is the absolute temperature and $l_0$ is the charge separation distance along a polymer chain. For $\Gamma < 1$ the entropic term is negligible and the complexation is driven by the change in the Coulomb energy, but for $\Gamma > 1$ the entropic term starts to play a more important role, for $\Gamma < 1.5$ the electrostatic attraction between the oppositely-charged polyelectrolytes still predominantly drives the complexation, while the counterion release entropy only plays a subsidiary role. For $\Gamma > 1.5$, the contribution of the entropic term becomes more significant, until at $\Gamma = 2.5$ the complexation is completely driven by counterion release entropy (13).

The charge separation distance $l_0$ is 0.76 nm (two times the bond length) for R-DPRs, and 0.38 nm (the bond length) for acidic molecules. Using the average $l_0 = (0.76 + 0.38)/2 = 0.57$ and $l_{\mathrm{B}} \approx 0.7$ nm for water at 300 K, the Coulomb interaction parameter is found to be $\Gamma = 1.23$ that lies in the range where the Coulomb energy change plays a more significant role than the counterion release entropy, supporting our assumption to neglect the effect of counterion condensation.

### Simulations

**Single-molecule simulations.** Langevin dynamics simulations are performed at 300 K in NVT ensembles with a time-step of 0.02 ps and a Langevin friction coefficient of 0.02 ps$^{-1}$ using GROMACS version 2016. Each simulation is performed for

DRAFT

3 $\mu$s and the last 1 $\mu$s is used to obtain the $R_\mathrm{h}$ and $R_\mathrm{g}$. The Hydro++ program (19) is used to obtain the $R_\mathrm{h}$ values from the trajectories. The fits for $R_\mathrm{h}$ in Fig. 1*B* are according to $bN^\nu$. For $R_\mathrm{g}$ we use the following equation for the fits in Fig. 1*C* (20):

$$R_\mathrm{g} = \sqrt{\frac{2l_\mathrm{p}^* b}{(2\nu+1)(2\nu+2)}}N^\nu,$$

where $b = 0.38$ nm and $l_\mathrm{p}^* = 0.40 \pm 0.07$ nm. The errors represent the changes in the scaling exponents for $0.33 < l_\mathrm{p}^* < 0.47$. The asphericity in Fig. 1*D* is calculated using the following equation (21):

$$\text{Asphericity} = 1 - 3\left\langle \frac{\lambda_1\lambda_2 + \lambda_2\lambda_3 + \lambda_3\lambda_1}{(\lambda_1+\lambda_2+\lambda_3)^2} \right\rangle,$$

where $\lambda_{1,2,3}$ are the eigenvalues of the gyration tensor. Asphericity is zero for a perfect sphere and is 1 for a perfect rod.

**Droplet simulations and cluster size distribution analysis.** To perform droplet simulations the DPRs are placed in a cubic box of size 80 nm and then simulated for $\approx 3$ $\mu$s which is sufficient to reach the equilibrium state (Figs. 2*A* left panels, 3*B*, S3-S7). At equilibrium the number of the residues inside the condensed phase (droplet) and the radial density profile measured from the center of the droplet (e.g. Fig. S5) are well-converged. In our test simulations we observed no significant effect of the initial distribution of the molecules on the properties of the resulting droplets at equilibrium. The radial density profiles (Figs. 3*C*, S5) and cluster size distribution plots (Figs. S3*A*, S4) are time-averaged for at least 1 $\mu$s at equilibrium. Discrete cells of thickness 1 nm are used to obtain the radial density profiles. The droplet radius is obtained from the radial density profile and is the average position of the first point close to the low-density region with $|d\rho/dr| \leq$ 2-5 mg/ml/nm (see dashed lines in Figs. 3*C*, S5). This term is the absolute value of the slope of the density profile which is close to zero where the density profile reaches the dilute phase region. Note that using $|d\rho/dr| \approx 0$ mg/ml/nm gives unrealistically large values for the droplet radius. We performed a sensitivity analysis for the critical value of $|d\rho/dr|$ and observed that selecting slope limits between 2-5 mg/ml/nm results in a maximal change of 4 % for all computed values for the droplet radius.

To generate the cluster size distribution plots, two chains are considered to be in the same cluster if at least two residues of those chains come closer than 0.7 nm (22). In our cluster size distribution plots (Fig. S3*A* and S4) the horizontal axis is the logarithm of the number of residues inside a cluster ($S$) and the vertical axis is the logarithm of the time-averaged number of the clusters ($N_\mathrm{C}$). When phase separation occurs the curves are divided into two regions, a dilute phase containing free molecules and small clusters, and a condensed phase that exchanges molecules with the dilute phase.

**Slab simulations.** The initial simulations are performed in cubic boxes of size 20 nm for poly-GA and 25 nm for more extended R-DPRs. These box sizes ensure no interaction between DPRs and their periodic images for repeat lengths $n \leq 100$. For longer DPRs the box size is increased accordingly. For the initial equilibration simulations we followed the steps suggested in (23). After equilibration, the box is enlarged to a 10 times larger size than its initial value in the $z$ direction which is sufficient to reduce the finite-size effects and to obtain reliable values for $\rho_\mathrm{H}$ and $\rho_\mathrm{L}$. The system is then simulated for $\approx 3$ $\mu$s in an NVT ensemble to achieve convergence for the density profiles in the $z$ direction (Figs. 2*B*, S10, S11*B*). The density profiles are calculated using discrete cells of thickness 1 nm and time-averaged for at least 1 $\mu$s at equilibrium. When the system undergoes phase separation, the averaged concentrations in $|z| < 4$ nm region is used to obtain $\rho_\mathrm{H}$. To obtain $\rho_\mathrm{L}$ we use the average concentrations in $|z| > 40$ nm for poly-GA (Fig. 2*B*) and $|z| > 65$ nm for R-DPRs (Figs. S10, S11*B*). The simulation parameters are similar to the ones we used in the single-molecule and droplet simulations.

**Phase diagrams.** The vertical axis in our phase diagram is the DPR repeat length and the horizontal axis is the concentration (Figs. 2*A*, 4*A*, S11*A*). The phase diagram is obtained by connecting values of $\rho_\mathrm{L}$ and $\rho_\mathrm{H}$ computed from the slab simulations. To find the critical point, we begin with the smallest repeat length $n_1$ that has produced converged $\rho_\mathrm{L}$ and $\rho_\mathrm{H}$ values. Then we perform slab simulation for repeat length of $n_{-1} = n_1 - 10$ to calculate the time-averaged density profile for 1 $\mu$s after 3 $\mu$s of simulation time. If the calculated density profile for repeat length $n_{-1}$ is almost flat, with small fluctuations in concentration $\Delta\rho = |\rho_\mathrm{max} - \rho_\mathrm{min}| < 20$ mg/ml, we report $\frac{n_1+n_{-1}}{2}$ as the critical repeat length. With this method we estimate the critical repeat length with an error of less than 5 repeats. Choosing any value larger than 20 mg/ml for $\Delta\rho$ does not change the critical repeat length. In each phase diagram, for the region close to the critical point, a dashed spline that reaches its minimum at $(\frac{n_1+n_{-1}}{2}, \frac{\rho_\mathrm{H}(n_1)+\rho_\mathrm{L}(n_1)}{2})$ is shown as a guide to the eye (Figs. 2*A*, 4*A*, S11*A*).

**Potential of mean force (PMF) calculation.** We use umbrella sampling simulations and the weighted histogram analysis method (WHAM) via the gmx wham utility of GROMACS to calculate the PMF associated with the binding of R-DPRs to the acidic molecules in Fig. S8. The distance $r$ between the center of masses of two molecules is considered as the reaction coordinate. For each window the simulation is conducted for 1 $\mu$s. We use $\Delta r = 0.1$ nm for the distance between two windows. For details about the umbrella sampling method the reader is referred to (3, 24).

### Protein sequences

Sequences of the disordered parts of NPM1 and NCL used in Fig. S9 are listed below. The negatively charged residues are shown in blue.
$\mathrm{NPM1}_{120-240}$

EEDAESEDEEEEDVKLLSISGKRSAPGGGSKVPQKKVKLAADEDDDDDDEEDDDEDDDDDDFDDEEAEEKAPVKKSIRDTPAKNAQKSNQN
GKDSKPSSTPRSKGQESFKKQEKTPKTPKG

$\mathrm{NCL}_{1-300}$

MVKLAKAGKNQGDPKKMAPPPKEVEEDSEDEEMSEDEEDDSSGEEVVIPQKKGKKAAATSAKKVVVSPTKKVAVATPAKKAAVTPGKKAAA
TPAKKTVTPAKAVTTPGKKGATPGKALVATPGKKGAAIPAKGAKNGKNAKKEDSDEEEDDDSEEDEEDDEDEDEDEDEIEPAAMKAAAAAPA
SEDEDDEDDEDDEDDDDDEEDDSEEEAMETTPAKGKKAAKVVPVKAKNVAEDEDEEEDDEDEDDDDDEDDEDDDDEDDEEEEEEEEEEPVK
EAPGKRKKEMAKQKAAPEAKKQKVEG

### Supporting figures

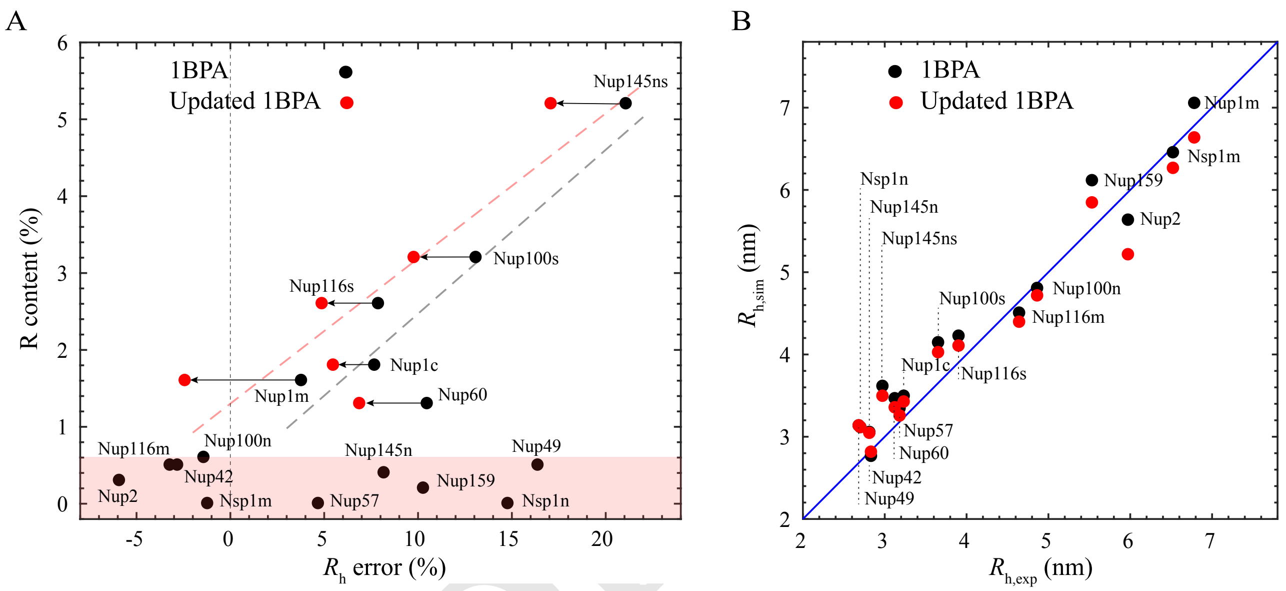


**Fig. S1.** (A) Content of Arginine in FG-Nup segments plotted against their corresponding hydrodynamics radius error: $(R_{\mathrm{h,sim}} - R_{\mathrm{h,exp}})/R_{\mathrm{h,exp}}$ for 1BPA and the updated 1BPA force fields. The red shaded band contains FG-Nups with R content $< 0.6$ %. The black dashed line shows the correlation between the R content $> 0.6$ % and the $R_{\mathrm{h}}$ error in 1BPA. In the updated 1BPA force field the $R_{\mathrm{h}}$ error is reduced for all FG-Nups with R content $> 0.6$ %. The red dashed line shows the correlation between the R content $> 0.6$ % and the $R_{\mathrm{h}}$ error in updated 1BPA. (B) A direct comparison of the two force fields in predicting the hydrodynamic radius of FG-Nups. The total average and the largest errors are found to be $8.3\%$ and $21.1\%$ in the 1BPA force field, and $7.5\%$ and $17.1\%$ in the updated 1BPA force field.

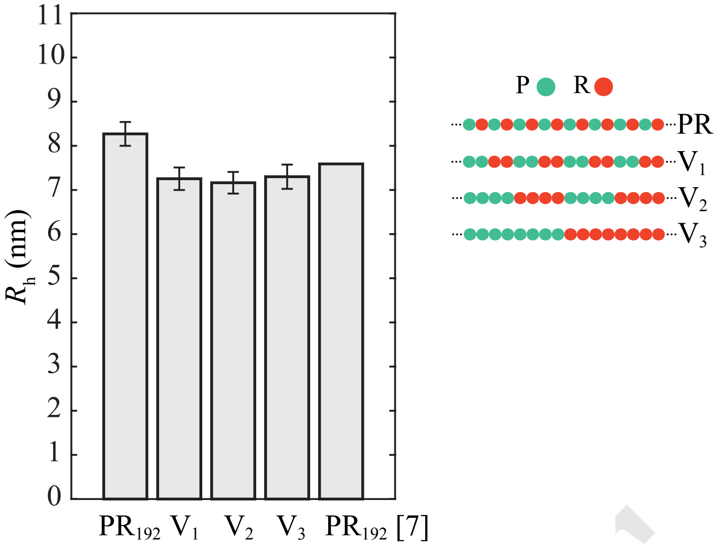


**Fig. S2.** The $R_\mathrm{h}$ value of poly-PR and three variants with different patterning of the Proline and Arginine residues. The bar chart at the right has been obtained using the equation reported in (7).

DRAFT

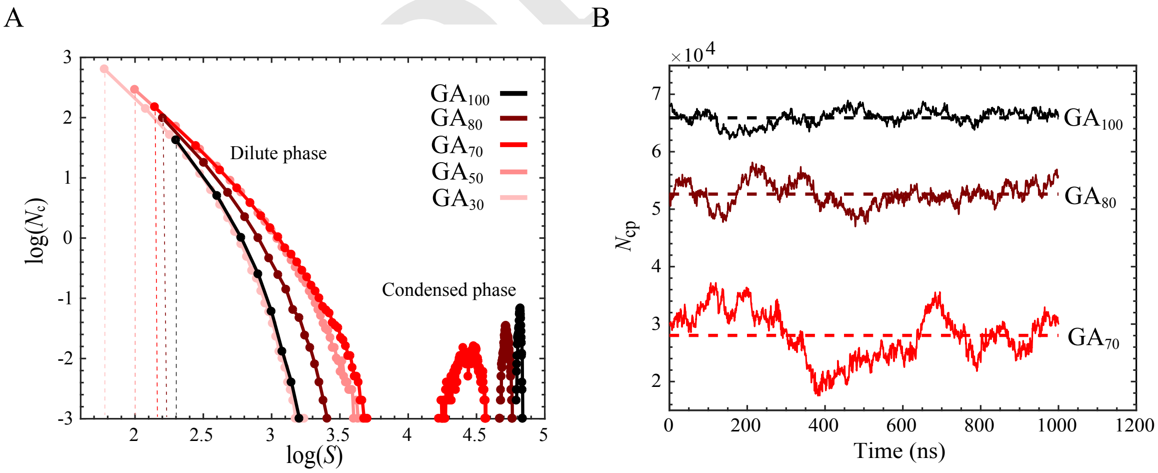


**Fig. S3.** Phase separation of poly-GA for a total concentration of 32.2 mg/ml. (A) Cluster size distribution of $GA_n$ $(30 \leq n \leq 100)$ at equilibrium. $S$ is the number of residues inside a cluster and $N_\mathrm{C}$ is the time-averaged number of the clusters. For comparison, the time-averaged number of free molecules is also included in this plot, indicated with a dashed line for each case. (B) The number of residues in the condensed phase $(N_\mathrm{cp})$ plotted against time at equilibrium for $GA_{100}$, $GA_{80}$, and $GA_{70}$. Horizontal dashed lines indicate the average values. Longer dipeptides form clusters containing more residues with lower exchange with the surrounding.

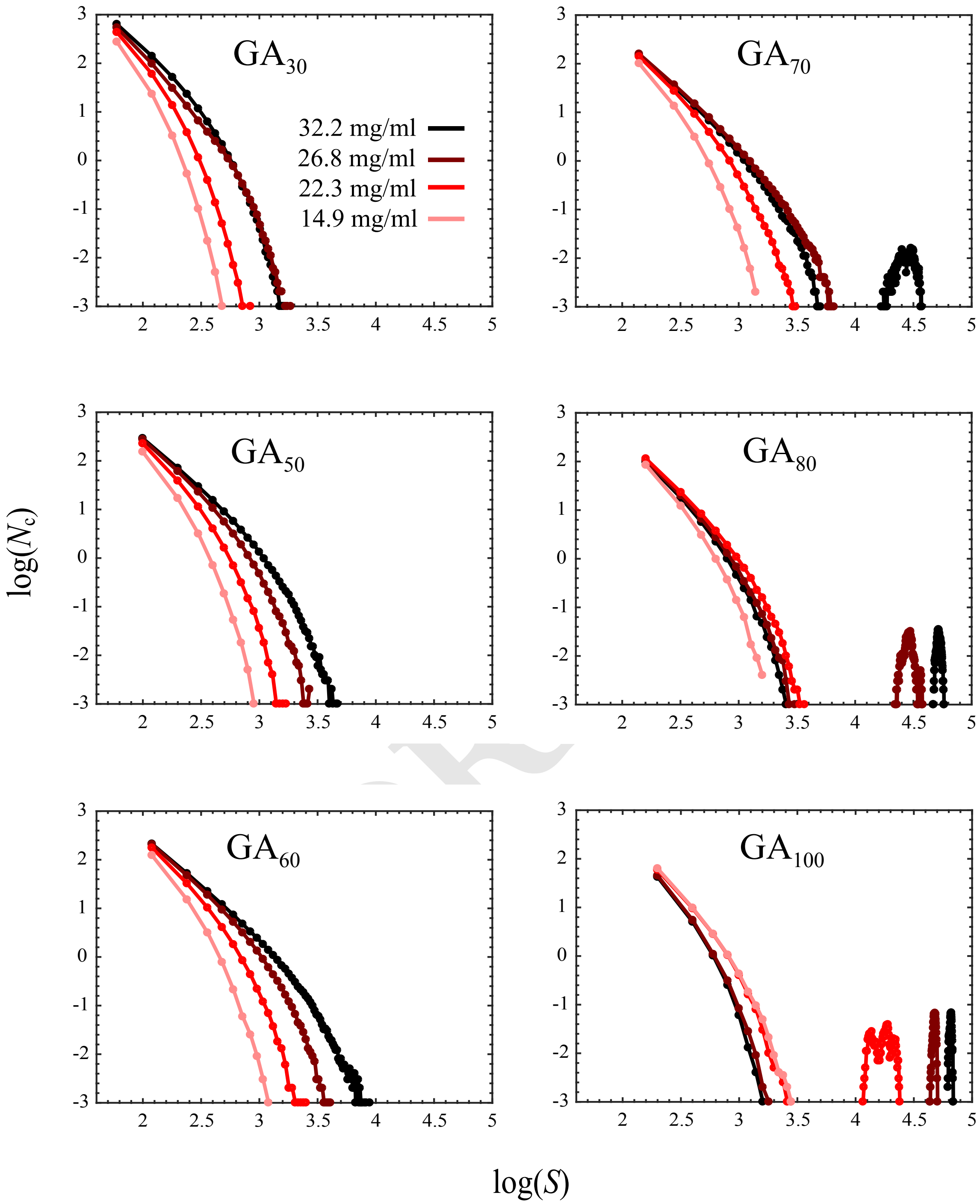


**Fig. S4.** Cluster size distribution analysis of poly-GA at equilibrium for four different total mass concentrations of 32.2, 26.8, 22.3, and 14.9 mg/ml shows a length- and concentration-dependent phase separation. $S$ is the number of residues inside a cluster and $N_\mathrm{c}$ is the time-averaged number of the clusters. For a fixed repeat length, increasing the concentration increases the average number of the molecules inside the condensed phase (see the results for $\mathrm{GA}_{80}$ and $\mathrm{GA}_{100}$).

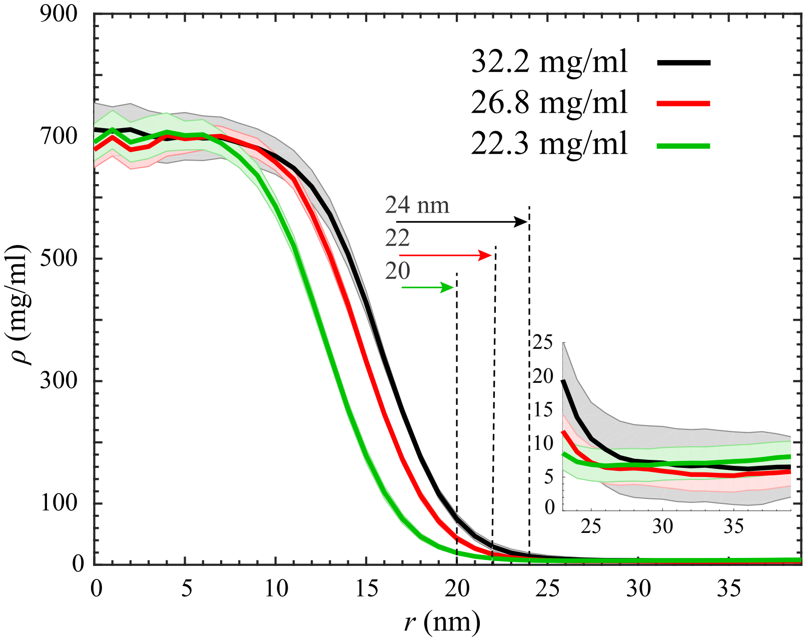


**Fig. S5.** Radial density profiles for $GA_{100}$ droplets for three different total mass concentrations. The shading indicate half of the standard deviations as error bars. The radius of the droplet is shown with dashed lines for each case. The inset figure shows the zoomed density profiles for $r \geq 24$ nm. The size of the droplet increases with increasing the total mass concentration. However, $\rho_{\mathrm{H}}$ (the average concentration for $r < 4$ nm) and $\rho_{\mathrm{L}}$ (the average concentration for $r > 30$ nm) remain unchanged.

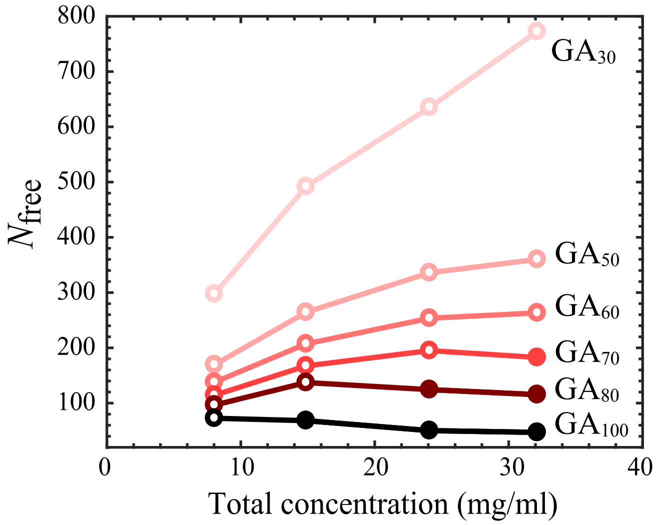


**Fig. S6.** Number of free molecules $N_{\mathrm{free}}$ in the dilute phase plotted against the total concentration for different lengths of poly-GA. Filled markers are used when poly-GA molecules undergo phase separation. When phase separation occurs the $N_{\mathrm{free}}$ drops. At a fixed concentration, $N_{\mathrm{free}}$ is higher for shorter dipeptides. The data in the figure is obtained from the cluster size distribution curves of Fig. S4.

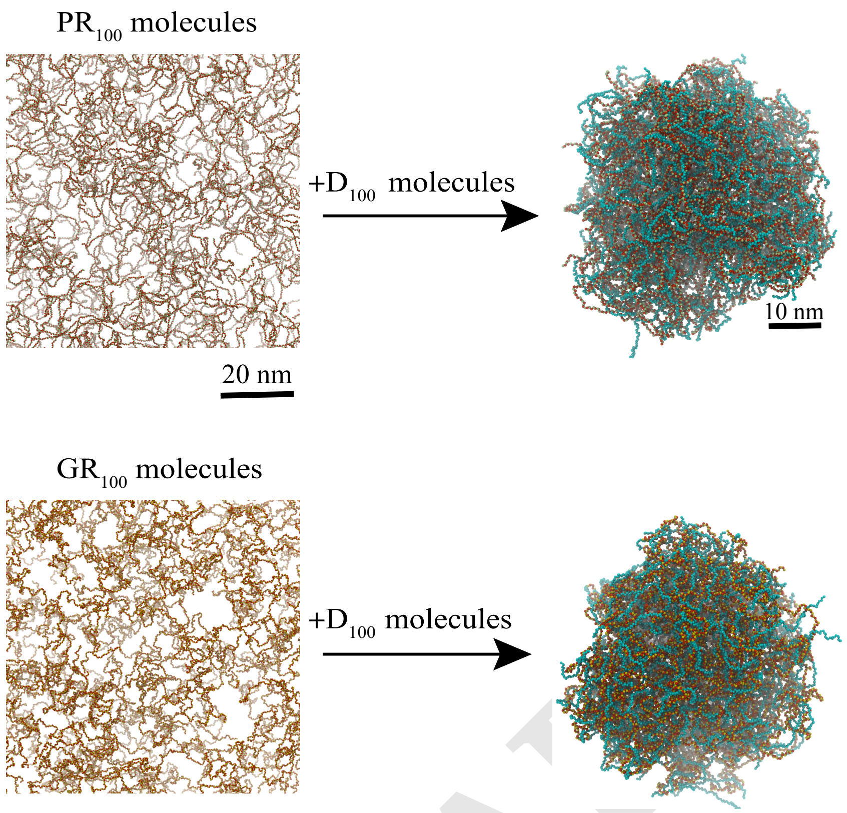


**Fig. S7.** Long chains of poly-PR and poly-GR are not capable of forming clusters (left). Adding acidic molecules (poly-D) induces the phase separation of R-DPRs (right) for poly-PR concentration ratio of $r_{\rm PR} = 0.57$. The total concentration is 14.8 mg/ml for all cases.

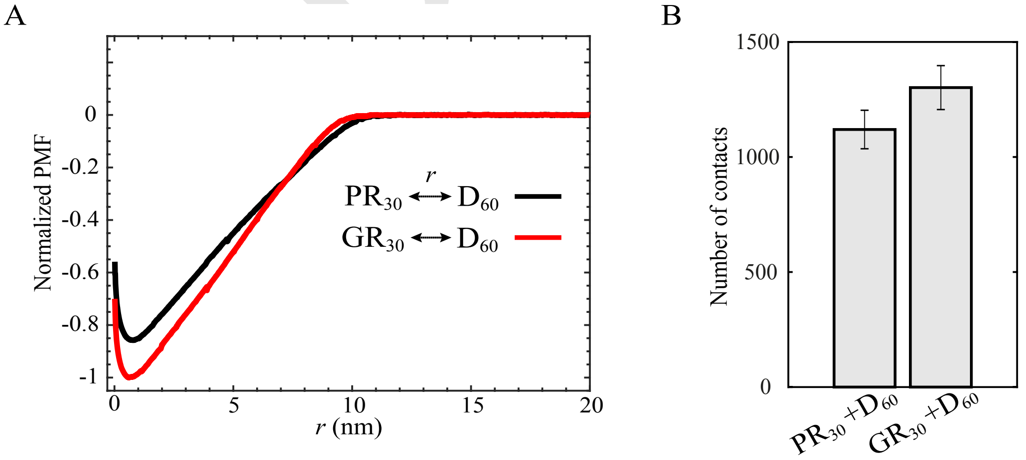


**Fig. S8.** (A) PMF curves for binding of PR$_{30}$ and GR$_{30}$ to D$_{60}$ indicates a larger free energy of binding of poly-GR to acidic molecules. The distance between the center of masses of two molecules is indicated with $r$. Due to the more compact conformation of poly-GR, its PMF curve vanishes at slightly shorter inter-molecule distances. Both curves are normalized with the depth of the GR$_{30}$-D$_{60}$ binding free energy. (B) Time-averaged total number of contacts after binding of PR$_{30}$ and GR$_{30}$ to D$_{60}$ at equilibrium using a cut-off of 2.5 nm. The contact of one residue in the R-DPRs with one residue in the acidic molecule is counted as one contact.

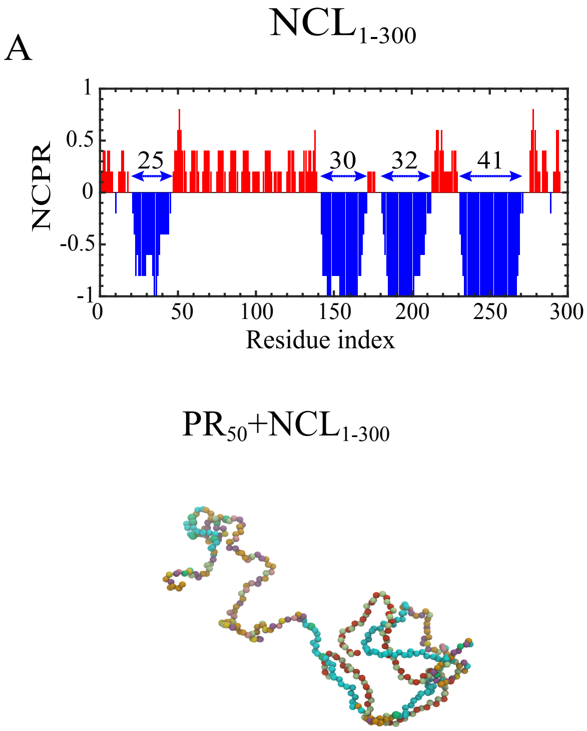


**Fig. S9.** Top: Net charge per residue (NCPR) histograms for disordered parts of two nucleolar proteins: (A) nucleolin ($NCL_{1-300}$) and (B) nucleophasmin ($NPM1_{120-240}$). Blue arrows show acidic tracts with lengths ranging from 12 to 41. To find NCPR we use a sliding window containing 5 residues. Bottom: snapshots of binding of PR50 to $NCL_{1-300}$ and $NPM1_{120-240}$. Acidic tracts are indicated in cyan; PR chains comprise of red-green colored beads (as in Fig. 1*A*). The other aminoacids are given a range of colors according to their aminoacid type.

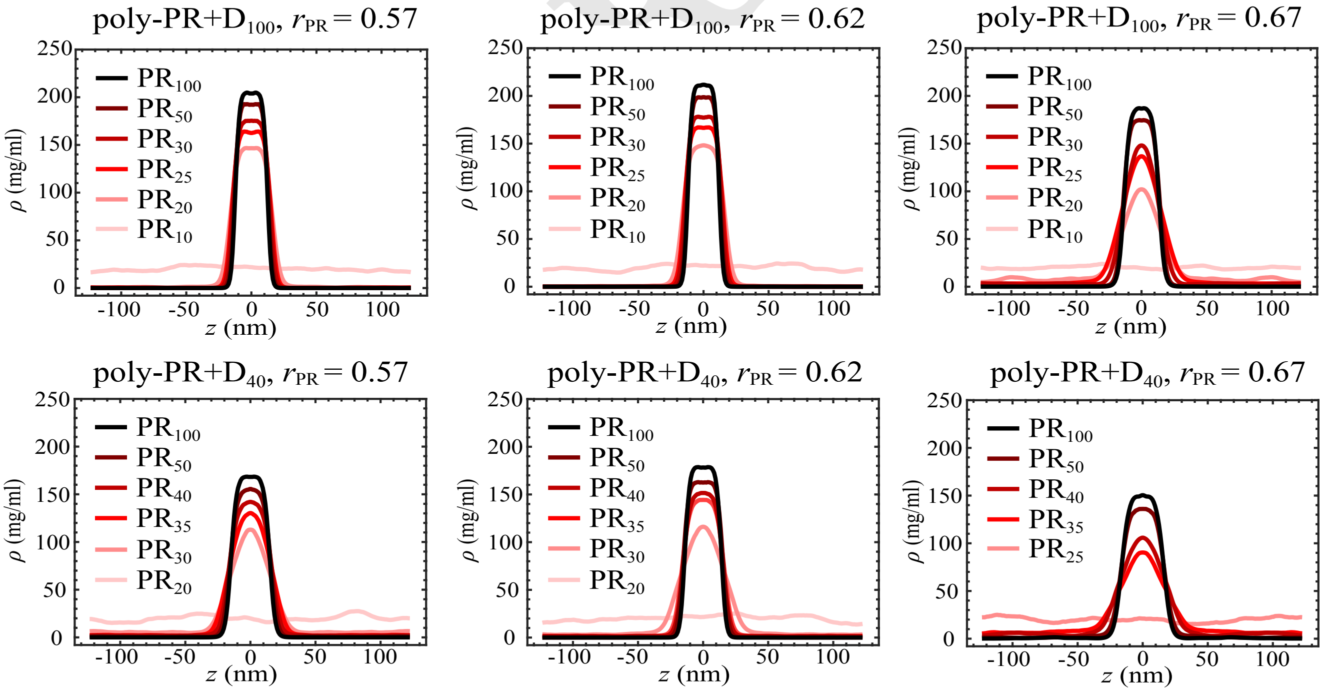


**Fig. S10.** Slab density profiles for phase separation of poly-PR with acidic molecules of lengths 40 and 100 for three different concentration ratios of $r_{PR}$ = 0.57, 0.62, and 0.67. These density profiles are used to obtain the coexistence phase diagrams in Fig. 4*A*.

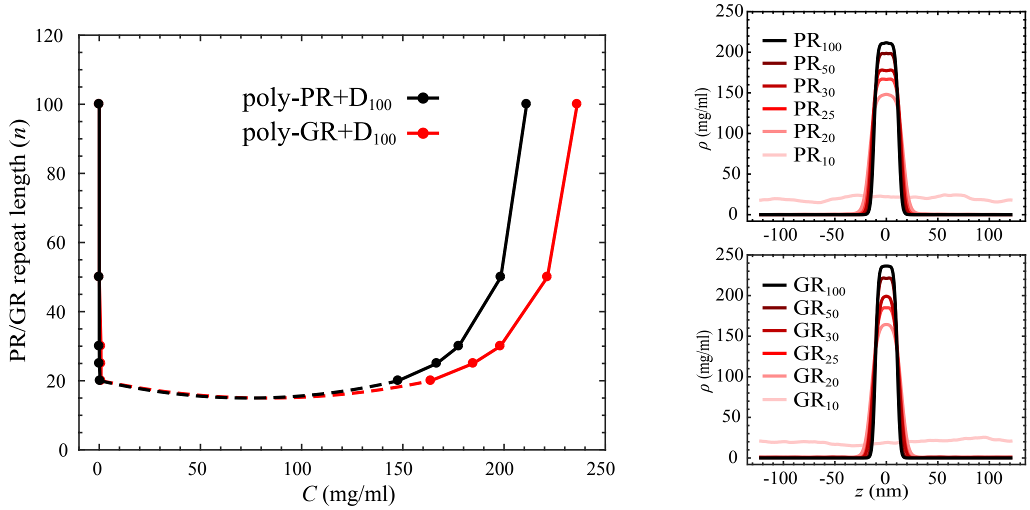


**Fig. S11.** Coexistence phase diagram (left) and the corresponding slab density profiles (right) for phase separation of poly-PR and poly-GR with $D_{100}$ for $r_{\mathrm{PR}} = r_{\mathrm{GR}} = 0.62$. With the same repeat lengths, poly-GR forms condensed phases with higher concentrations.

### Supporting tables

**Table S1.** The relative hydrophobic strength $\varepsilon_i$ values of charged residues in the 1BPA force field (1, 2) and updated 1BPA force field (the current paper)

| Amino acid | R | D | E | K |
|---|---|---|---|---|
| $\varepsilon_i$ (1BPA) | 0 | 0.0005 | 0.0005 | 0.0005 |
| $\varepsilon_i$ (updated 1BPA) | 0.005 | 0.005 | 0.005 | 0.005 |

DRAFT
